# Interpreting Generative Adversarial Networks to Infer Natural Selection from Genetic Data

**DOI:** 10.1101/2023.03.07.531546

**Authors:** Rebecca Riley, Iain Mathieson, Sara Mathieson

## Abstract

Understanding natural selection in humans and other species is a major focus for the use of machine learning in population genetics. Existing methods rely on computationally intensive simulated training data. Unlike efficient neutral coalescent simulations for demographic inference, realistic simulations of selection typically requires slow forward simulations. Because there are many possible modes of selection, a high dimensional parameter space must be explored, with no guarantee that the simulated models are close to the real processes. Mismatches between simulated training data and real test data can lead to incorrect inference. Finally, it is difficult to interpret trained neural networks, leading to a lack of understanding about what features contribute to classification.

Here we develop a new approach to detect selection that requires relatively few selection simulations during training. We use a Generative Adversarial Network (GAN) trained to simulate realistic neutral data. The resulting GAN consists of a generator (fitted demographic model) and a discriminator (convolutional neural network). For a genomic region, the discriminator predicts whether it is “real” or “fake” in the sense that it could have been simulated by the generator. As the “real” training data includes regions that experienced selection and the generator cannot produce such regions, regions with a high probability of being real are likely to have experienced selection. To further incentivize this behavior, we “fine-tune” the discriminator with a small number of selection simulations. We show that this approach has high power to detect selection in simulations, and that it finds regions under selection identified by state-of-the art population genetic methods in three human populations. Finally, we show how to interpret the trained networks by clustering hidden units of the discriminator based on their correlation patterns with known summary statistics. In summary, our approach is a novel, efficient, and powerful way to use machine learning to detect natural selection.

## 1 Introduction

In the last several years, numerous deep learning methods have been proposed as solutions to population genetic problems (see [1] for a recent review), in part due to the ability of deep neural network architectures to recover evolutionary signals from noisy population-level data. Convolutional neural networks (CNNs) have been particularly effective and allowed the field to move away from summary statistics and towards analyzing haplotype matrices directly. The broad idea behind CNNs for genetic data is to treat these matrices (with individuals/haplotypes on the rows and sites/SNPs on the columns) as analogous to images, with convolutional filters picking up on correlations between nearby sites. Population genetic CNNs have now been developed for a variety of applications, including recombination hotspot identification [2], introgression, recombination rates, selection, and demography [3], natural selection [4, 5], adaptive admixture and introgression [6, 7], balancing selection [8], dispersal inference [9], and the task-agnostic software `dnadna` [10]. New architectures (including those built upon CNNs) have also been proposed, including graph neural networks (GNNs) [11], and U-Nets [12], and recurrent neural networks (RNNs) [13, 14].

One drawback of these machine learning approaches is that they require simulated training data, since labeled examples (i.e. where the historical evolutionary “ground truth” is known) are limited. If the simulated training data is not a good match for the real genetic data, inference results may not be robust. This is a particular problem for understudied populations and non-model species, where less is known about their evolutionary parameter space. Several solutions to this “simulation mis-specification” problem have been proposed, including adaptive re-weighting of training examples [15] and domain-adaptive neural networks [16]. Another strategy is to create custom simulations using Generative Adversarial Networks (GANs) [17], which have recently been developed for population genetic data. Our previous work [18] created a GAN (`pg-gan`) that fits an evolutionary model to any population – as training progresses, the generator produces synthetic data that is closer and closer to the real data, confusing the discriminator. The key innovation of `pg-gan` is that it learns an explicit evolutionary generative model, in contrast to other GANs which generate sequences that look like real data from random processes with no underlying model [19, 20]. Recent work has improved GAN-style approaches for population genetics using adversarial Monte Carlo methods [21].

GANs have two components, a generator which creates synthetic examples, and a discriminator which classifies examples as real or fake. After training, the focus is often on the generator, which can be used to produce novel results (i.e. faces, text, genetic data) from a vector of random noise. In `pg-gan` the generator is not a neural network but instead an evolutionary model. The fitted generator from `pg-gan` is therefore useful in its own right, as it is directly interpretable in terms of population genetic parameters. Because the specification of such models is relatively standardized (population size changes, splits, migrations, admixtures, etc), we can easily use the resulting model in downstream applications or make it available to other researchers in a standard format (see [22]). For example, we could use the demography as a null model or as training or validation data for a second machine learning method.

Although well-trained discriminators can be used downstream to validate the examples created by other generators (e.g. [23]), there is typically less emphasis on using or analyzing the trained discriminator. In this work we develop a new approach that uses the trained discriminators of `pg-gan` to identify non-neutral regions. The major advantage of this approach over existing approaches is that it requires very few simulated selected regions during training. Despite recent advances (e.g. [24]), simulating realistic natural selection is very difficult, with many parameter choices that make mirroring real data challenging. These include the type of selection, time of onset of selection, selection strength, and frequency of the selected variant at the present. For models with exponential growth or large population sizes, forward simulations become prohibitively computationally expensive. Our approach bypasses the need to simulate large numbers of selected regions by instead detecting regions of real data that do not conform to an inferred (neutral) demographic model. This approach is similar to traditional population genetic approaches to detecting selection by looking for regions that are outliers in terms of divergence, diversity or haplotype structure [25]. However instead of looking at one or a combination of summary statistics [26], our approach operates directly on the observations, and in principle is able to use all the information present in the data.

Understanding what neural networks actually learn has been a major goal of interpretability work for many years. Despite progress in the image domain (e.g. [27–29], less is known about what CNNs for population genetic data are learning (see [30] for recent progress). Our interpretability analysis links traditional summary statistics (which are known to be informative for selection and other evolutionary parameters) with the hidden nodes of the discriminator network. This gives us a window into the network’s ability to compute existing summary statistics, importance and redundancy of different statistics, and differences between discriminators.

Overall our method represents a novel approach to detect natural selection and other types of non-neutrality, while avoiding both demographic parameter mis-specification and extensive selection simulations. Our software is available open-source at https://github.com/mathiesonlab/disc-pg-gan.

## 2 Materials and Methods

### 2.1 GAN training

Our existing GAN application, `pg-gan`, consists of a CNN-based discriminator and an `msprime`-based generator that are trained in concert. During training, the generator selects parameters to use for the simulation software `msprime` [31, 32], creating “fake” regions. The discriminator learns to distinguish these simulated neutral regions from the real training data, which contains both neutral and selected regions. With feedback from the discriminator, the generator learns which parameters will produce the most realistic simulated data, and the discriminator gets better at picking up on differences between the two types of data. `pg-gan` is implemented in `tensorflow` [33] and can be run on a GPU using `CUDA` [34]. As the discriminator is a CNN, we are able to train it using traditional gradient descent approaches (i.e. backpropagation). However, since we currently cannot pass gradients through the coalescent with recombination, we train the generator using a gradient-free approach (in this case simulated annealing, but other methods are possible). At the end of this optimization process, we obtain a generator that is good at creating realistic simulated data, and a discriminator that can identify real data accurately [18].

In this work we focus on the trained discriminator rather than the generator. We modify `pg-gan` to save the trained discriminator network. The input to this CNN is a genomic region represented by a matrix of shape (*n, S*, 2) where *n* is the number of haplotypes, *S* is the fixed number of consecutive SNPs, and there are two separate channels: one for the haplotype data and one for the inter-SNP distances (duplicated down the rows). These two channels are analogous to the three RGB color channels for images. For one-population evolutionary models, this network has two convolutional layers (32 and 64 filters each of dimension 1 *×* 5), two fully connected layers (each with 128 units), and an output of a single scalar value (probability of the example being real). We use binary cross-entropy as the loss function, with Adam optimization. With 1 *×* 2 max-pooling after each convolutional layer and a permutation invariant function before the fully connected layers, this results in 76,577 trainable parameters, all of which are saved after training.

### 2.2 Discriminator evaluation

In a small number of cases, GAN training (which is often subject to difficulties due to the minimax nature of the optimization problem) fails. In all our experiments, this manifested as discriminators predicting the same value for all regions (i.e. the network did not learn anything). We therefore discard these discriminators from further analysis.

### 2.3 Discriminator fine-tuning

Since the real data, unlike the simulated data, include regions that have experienced selection, the intuition is that regions that the discriminator predicts to be real with high probability correspond to non-neutral regions, since those have characteristics found in real but not simulated data. One concern is that there are processes other than selection that are found in the real but not simulated data (heterogeneity in parameter values, genotype or reference errors and so on) which might be identified by the discriminator. We therefore incentivize the discriminator to focus on selection in particular by *fine-tuning* it using a small number of forward simulations incorporating selection (see Section 2.6 below for details). Specifically, we use 3000 neutral regions and 2400 selected regions (600 each of selection strengths *s* = 0.01, 0.025, 0.05, 0.1). 20% of these regions are reserved for validation, and the discriminator is training on the remaining 80% for 1000 mini-batches, each of size 50 (roughly 12 epochs). We again use binary cross-entropy as the loss function, except now a label of 0 corresponds to neutral regions and a label of 1 corresponds to selected regions. This procedure *fine-tunes* the weights of the network such high discriminator predictions can be more reliably interpreted as evidence for selection.

### 2.4 Prediction

The fine-tuned discriminator can then be used to predict outcomes for new genetic regions. Values closer to 1 indicate more similarity to the real training data, and values closer to 0 indicate more similarity to the simulated data. A schematic of the entire workflow can be found in Figure 1.

**Figure 1.**
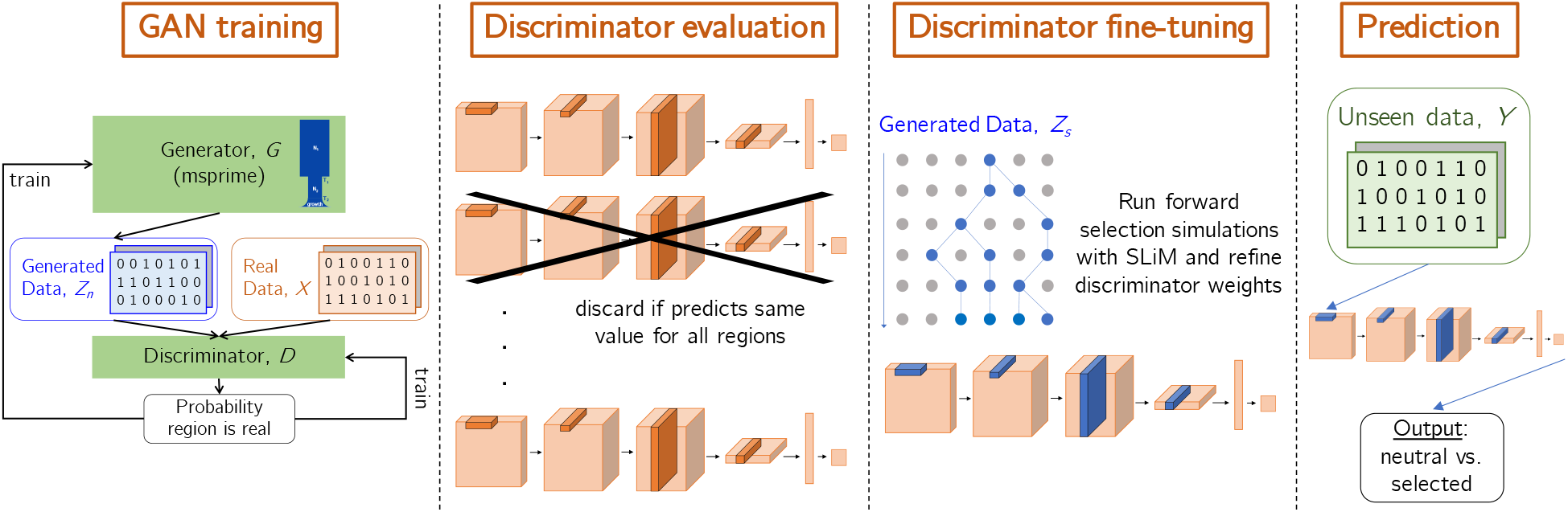
Selection inference workflow. First we run `pg-gan` many times (i.e. 20) to obtain a collection of discriminators. In the evaluation step, we discard any discriminator that does not pass quality checks (i.e. predicting the same value for every region). Retained discriminators (roughly 85%) are fine-tuned with a modest number of selection simulations. Finally, real data from a new but similar population is fed through each fine-tuned discriminator to obtain a probability. Regions where a large fraction of the discriminators produced a high probability are candidate selected regions.

We use ROC curves to evaluate the improvement in performance created by fine-tuning. Notably, in 9/10 cases of failed training, fine-tuning also fails to improve the discriminator – prediction results are the same before and after. In one case, a failed discriminator’s predictions (on selection simulations) were improved from fine-tuning, but we still discard it because it was not able to learn to generate realistic simulations initially. Below we describe additional method details and validation approaches, as well as an ensemble approach for successfully trained discriminators.

### 2.5 Population data and demographic models

To ensure that we are training and testing on different datasets, we use three pairs of similar populations from the 1000 Genomes Project [35]: CEU and GBR (Northern European), CHB and CHS (East Asian), and YRI and ESN (West African). These pairs were selected since they are the most similar in terms of genetic distance (see [35], Figure 2). Specifically, we train discriminators using CEU, CHB, and YRI then test on GBR, CHS, and ESN respectively. Although the number of samples may differ between the populations, this is not a problem due to the permutation-invariant architecture of the discriminator. Population samples sizes and number of regions are given in Table 1.

**Table 1:** Populations analyzed along with the number of individuals in each population. The number of regions is determined using non-overlapping 36-SNP windows, where at least 50% of bases must be in the accessibility mask.

| Code | Population description | # individuals | # regions |
| --- | --- | --- | --- |
| CEU | Utah residents with Northern and Western European ancestry | 99 | 292,936 |
| GBR | British in England and Scotland | 91 | 277,038 |
| CHB | Han Chinese in Beijing, China | 103 | 280,215 |
| CHS | Han Chinese South | 105 | 270,351 |
| YRI | Yoruba in Ibadan, Nigeria | 108 | 473,493 |
| ESN | Esan in Nigeria | 99 | 460,180 |

**Figure 2.**
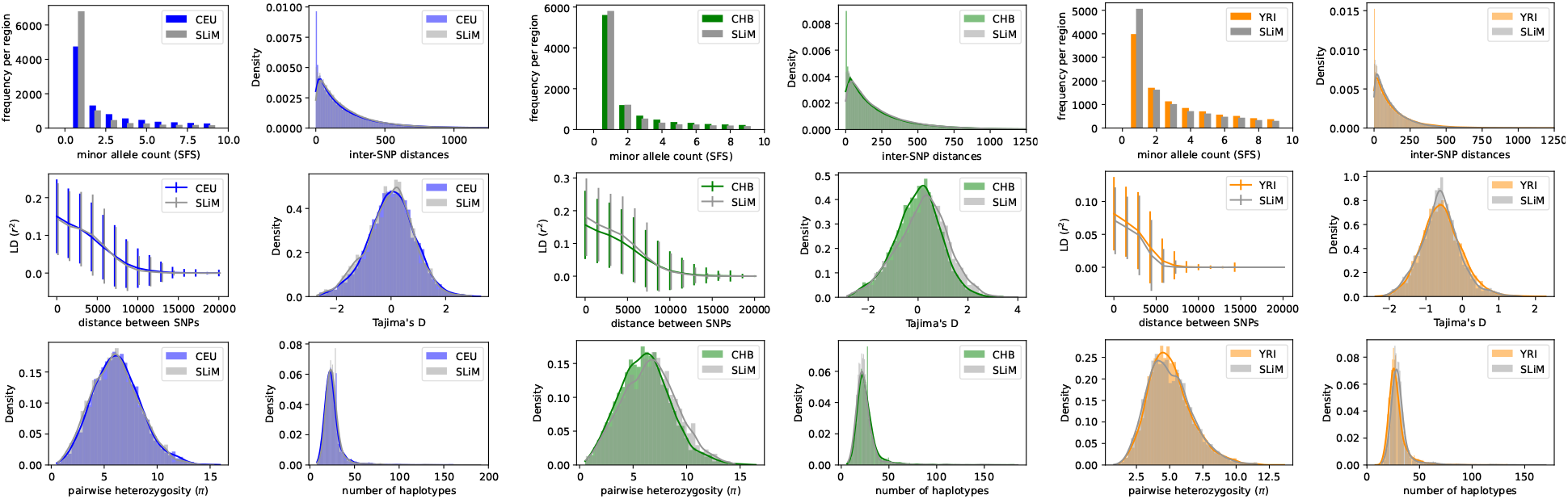
Summary statistic comparison of real training data (CEU, CHB, and YRI) to data simulated under `SLiM` with parameters inferred by `pg-gan`. The number of regions used for comparison was 5000.

For all populations we used an exponential growth model and fitted five parameters using `pg-gan`: the ancestral population size (*N*_1_), the timing of a bottleneck (*T*_1_), the bottleneck population size (*N*_2_), the onset of exponential growth (*T*_2_), and the growth rate (*g*). For all simulations we used a constant mutation rate of *µ* = 1.25 *×* 10^*−*8^ per base per generation and recombination rates drawn from the HapMap recombination map [36]. We simulated *L* = 50kb regions and then extracted the middle *S* = 36 SNPs. To avoid assuming an ancestral state, for each SNP we encode the minor allele as 1 and the major allele as *−*1. We found that centering the data around zero worked best with typical neural network activation functions, and also allowed us to zero-pad a small number of regions with fewer than *S* SNPs. For real data we filter non-biallelic SNPs and ensure that at least 50% of the bases are inside callable regions (see [35]), so that the discriminator avoids making real vs. fake distinctions based on artifacts.

### 2.6 Selection simulations for fine-tuning

To create the selection simulations used for fine-tuning, we use `SLiM` [24, 37], which can model a variety of types of natural selection. For the demography we use the same evolutionary model (exponential growth) as we used during `pg-gan` training. For European populations we use the parameters shown in Table 2, which were obtained from a run of `pg-gan` with CEU training data. During this training run, we capped the exponential growth parameter at 0.01, as allowing this parameter to be too large led to `SLiM` computations being too slow. See Figure 2 for a visualization of how well this simulated data matches CEU. Under this combination of parameters, we produced neutral regions, as well as regions with a single mutation undergoing positive selection with selection coefficients of of 0.01, 0.025, 0.05, and 0.1. For each selection coefficient we simulated 100 regions and recorded how many of them were classified as “fake”.

**Table 2:** Parameters of the exponential growth model, inferred for each population through a training run of `pg-gan`. There parameters are used for the demography in our `SLiM` selection simulations.

| population | $N_1$ | $N_2$ | $g$ | $T_1$ | $T_2$ |
| --- | --- | --- | --- | --- | --- |
| CEU | 22552 | 3313 | 0.00535 | 3589 | 1050 |
| CHB | 24609 | 3481 | 0.00404 | 4417 | 1024 |
| YRI | 23231 | 29962 | 0.00531 | 4870 | 581 |

We used a similar procedure to obtain a neutral demographic model for CHB and YRI. Due to the effect of large population sizes on `SLiM` runtimes, we again capped *g* at 0.01, and additionally capped *T*_2_ (the onset time of exponential growth) at 750 generations for YRI. The resulting parameters are shown in Table 2 and a visualization of the match is shown in Figure 2.

Selection is introduced by adding the mutation to one individual in the population, 1000 generations before the present. If the mutation is lost in any generation, we restart the simulation from the previous generation until we have reached the present – the resulting tree sequence can then be stored for further processing. First, the tree sequence is recapitated using coalescent simulations for efficiency (using the `pyslim` module). This step ensures that we have a complete tree sequence with all necessary ancestral nodes. Next, we sample the population down to *n* = 198 haplotypes, to match the real data. This allows us to prune and simplify the tree sequence before adding neutral mutations with `msprime`. For each region we store arrays with *S* SNPs, as well as arrays with all the SNPs, in order to compute Tajima’s D.

### 2.7 Validation on known selected regions

To validate predictions after fine-tuning, we examine each discriminator’s performance on genomic regions previously identified as targets of selection by Grossman *et al* [26] using a combination of different summary statistics. We use the 153 regions from their Table S1, including genes involved in metabolism, disease resistance, and skin pigmentation. We converted the start and end positions of each region to hg19 coordinates and sorted by population (CEU, CHB/JPT, and YRI). This resulted in a comparison between three types of data:

- Regions simulated using `msprime` under the demographic parameters inferred by `pg-gan` (corresponding to the current discriminator)
- Regions under positive selection as identified in [26]
- All other real regions (mostly neutral)

### 2.8 Discriminator ensemble method to detect selected regions

To assess the discriminator’s predictions genome-wide, we iterate through the entire genome (using non-overlapping 36-SNP windows) and make a prediction (probability of selection) for each region, then smooth the results by averaging probabilities in five consecutive windows. To visualize the results, we create modified Manhattan plots by plotting the probabilities on the y-axis (on a log scale). Individual discriminators can vary in their performance, so we also created an ensemble classifier by averaging predictions for each region over all successfully trained discriminators.

#### Algorithm 1: Ensemble method for using pg-gan discriminators to detect non-neutral regions.

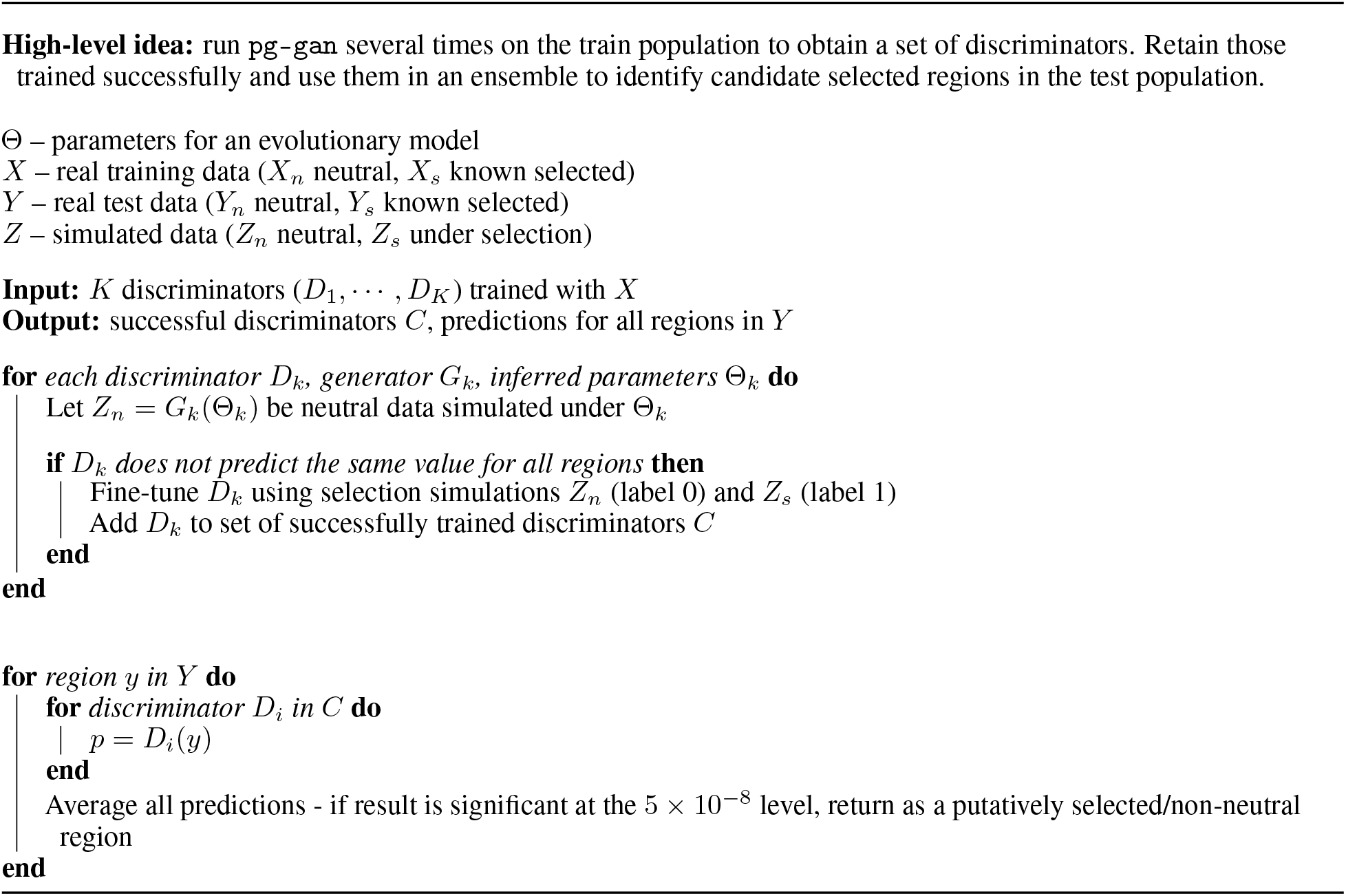

### 2.9 Interpretability analysis

Finally, to understand what the discriminator network is learning, we investigated whether the network had learned how to compute summary statistics that are known to be informative for evolutionary parameters (demography, selection, etc). To this end, we computed the correlation between the node values of the last hidden layer and various summary statistics. The motivation for analyzing the last hidden layer is that deep into the network, information from the haplotypes should be greatly distilled and processed, with informative high-level “features” extracted. The remaining step in the network is a linear combination of these hidden node values to create a logit value, which can be converted into a probability. We computed pairwise correlations between 128 hidden values and the following 61 summary statistics:

- The first 9 entries of the folded site frequency spectrum (SFS), excluding non-segregating sites.
- The 35 inter-SNP distances between the 36 SNPs of each region. These distances are rescaled to fall between 0 and 1.
- 15 linkage disequilibrium (LD) statistics (see [18] for details of the computation).
- Pairwise heterozygosity (*π*).
- The number of unique haplotypes across the 36 SNPs.

Note that the inter-SNP distances are also an input to the network. We calculated Pearson correlation coefficients and plotted them in a heatmap with summary statistics along rows and hidden values along the columns. To better visualize relationships between the hidden units, we clustered the columns according to their similarity using agglomerative clustering. The final visualization groups hidden nodes that are performing similar computations – since we perform a dropout (with rate 0.5) after each fully connected layer, we expect some redundancy in node behavior. We experimented with reducing the number of hidden units in the last few layers, as well as adding three fully connected layers (instead of two). However, all these modifications to the discriminator CNN resulted in degenerate training results (e.g. predicting all regions as real) so we pursued only the original architecture.

## 3 Results

### 3.1 Summary of discriminator training

For each training population (CEU, CHB, and YRI) we ran `pg-gan` 20 times to obtain a set of 60 discriminators. For each run we used a different seed, although even runs with the same seed can produce different results depending on the version of `tensorflow` and `CUDA`. In ten cases, training failed and the resulting discriminator predicted the same value for all regions (in other words, it ignores the input data – see Supplementary Figures S1 and S2 for an example). After we discarded these discriminators we were left with 50 (17 for CEU, 18 for CHB, and 15 for YRI). Below we discuss the outcomes of fine-tuning and prediction with three representative discriminators (one for each population pair), then explain how we used all successfully trained discriminators in an ensemble method to predict natural selection.

#### 3.1.1 Fine-tuning and validation on simulated data

In (A) of Figures 3, 4, and 5 we show the impact of fine-tuning on individual discriminators (seed 19 for CEU, seed 9 for CHB, and seed 4 for YRI). In the ROC curves for simulated data, a *positive* represents a region simulated under any strength of selection, and a *negative* represents a neutral region. For *real* neutral vs. selected data, we use the regions identified in [26] as the truth set. Dashed lines represent discriminator predictions before fine-tuning and solid lines represent predictions after fine-tuning. Fine-tuning improved discriminator predictions in all cases, although in a few cases the improvement was marginal. For YRI-trained discriminators tested on ESN, improvements were generally more marginal than for CEU- and CHB-trained discriminators.

**Figure 3.**
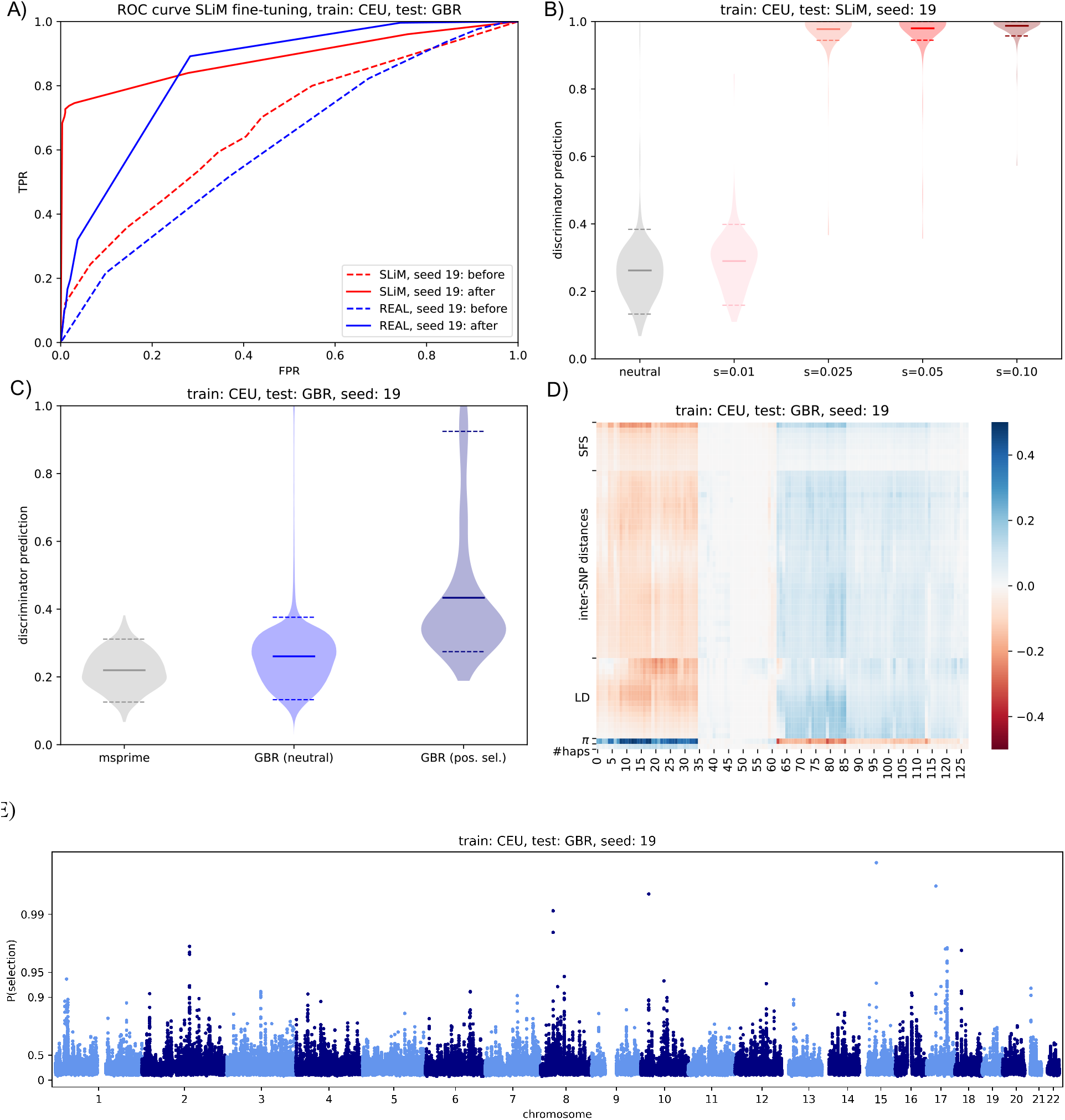
Example of a CEU-trained discriminator with test data from GBR (seed 19). A) ROC curve showing performance on selected regions before and after fine-tuning. All predictions (both simulated and real) are made for unseen (test) data. SLiM indicates simulated data with *s* = *xx* and REAL indicates selected regions from [26]. B) Performance of discriminator on unseen simulated data under various selection strengths. C) Performance of discriminator on selected regions from [26]. D) Correlation heatmap between discriminator hidden units (x-axis) and classical population genetics summary statistics (y-axis). The columns (hidden units) were clustered according to their similarity in terms of summary statistic correlation profiles. E) Genome-wide Manhattan plots of discriminator predictions on real test data.

**Figure 4.**
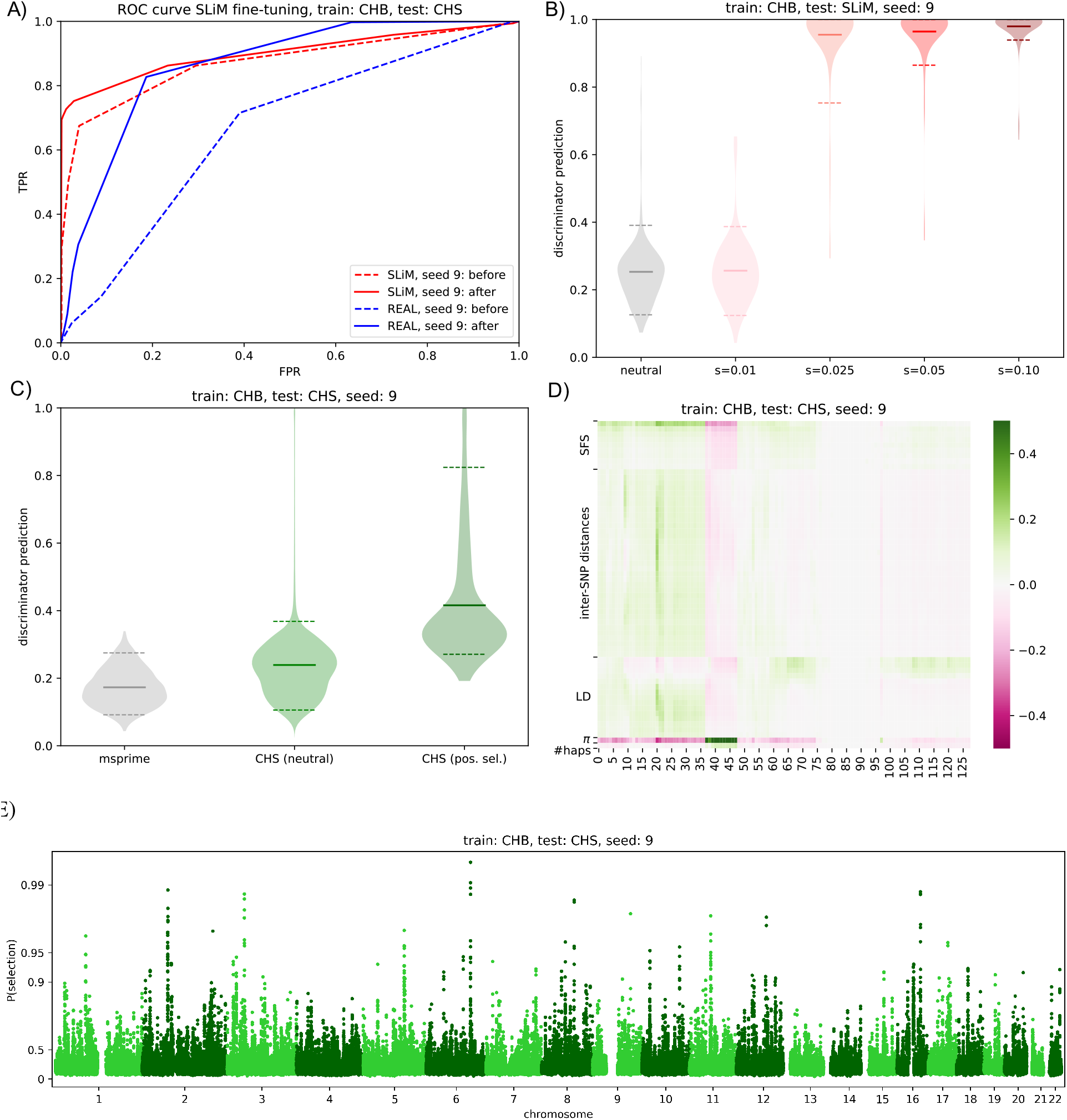
Example of a CHB-trained discriminator with test data from CHS (seed 9). A) ROC curve showing performance on selected regions before and after fine-tuning. All predictions (both simulated and real) are made for unseen (test) data. SLiM indicates simulated data with *s* = *xx* and REAL indicates selected regions from [26]. B) Performance of discriminator on unseen simulated data under various selection strengths. C) Performance of discriminator on selected regions from [26]. D) Correlation heatmap between discriminator hidden units (x-axis) and classical population genetics summary statistics (y-axis). The columns (hidden units) were clustered according to their similarity in terms of summary statistic correlation profiles. E) Genome-wide Manhattan plots of discriminator predictions on real test data.

**Figure 5.**
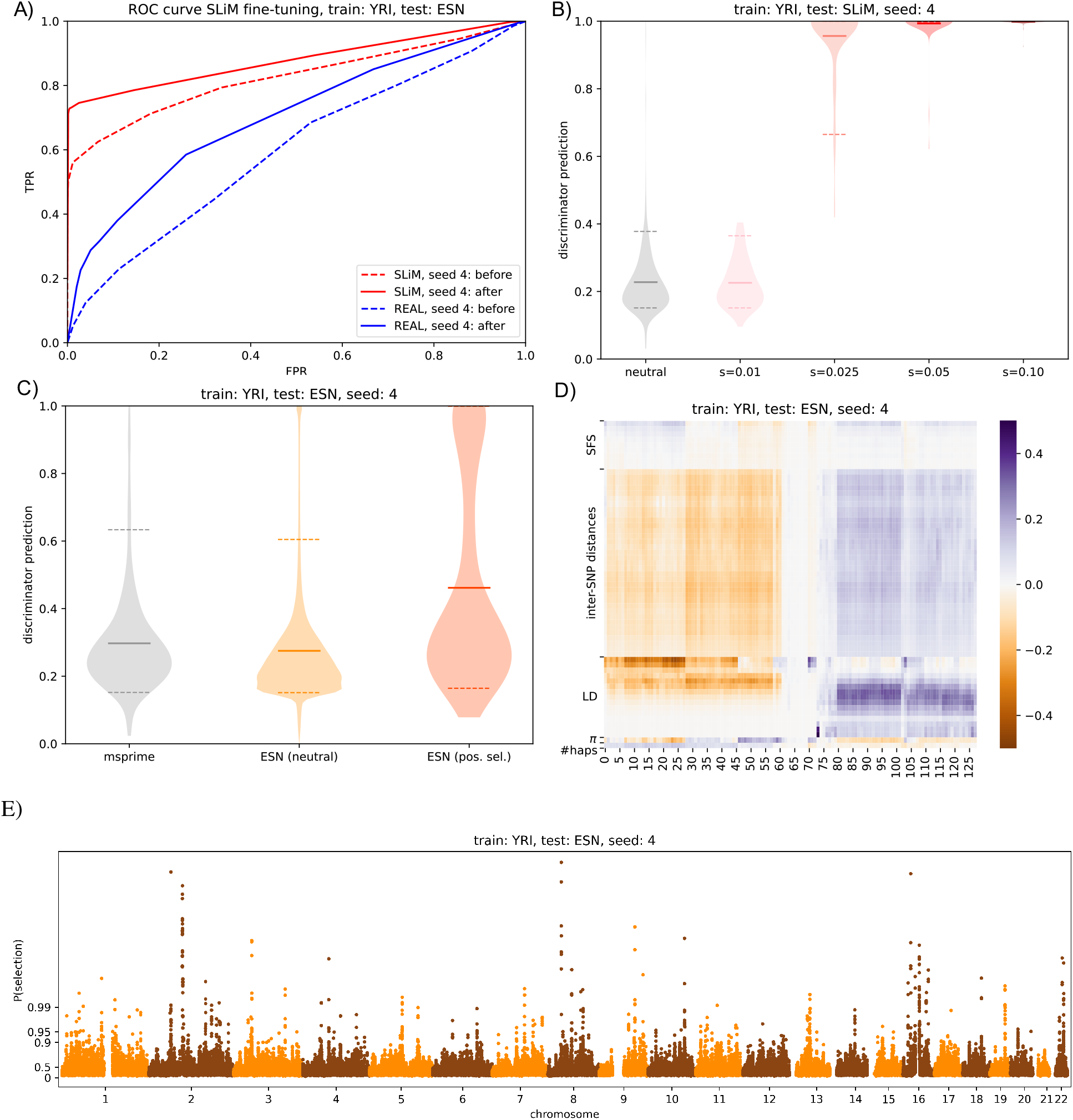
Example of a YRI-trained discriminator with test data from ESN (seed 4). A) ROC curve showing performance on selected regions before and after fine-tuning. All predictions (both simulated and real) are made for unseen (test) data. SLiM indicates simulated data with *s* = *xx* and REAL indicates selected regions from [26]. B) Performance of discriminator on unseen simulated data under various selection strengths. C) Performance of discriminator on selected regions from [26]. D) Correlation heatmap between discriminator hidden units (x-axis) and classical population genetics summary statistics (y-axis). The columns (hidden units) were clustered according to their similarity in terms of summary statistic correlation profiles. E) Genome-wide Manhattan plots of discriminator predictions on real test data.

After fine-tuning we run selection simulations (again created with `SLiM`) through each discriminator. In (B) of Figures 3, 4, and 5, we visualize these results through violin plots with 1000 neutral regions and 100 regions of each selection strength (all unseen during fine-tuning). Most discriminators group neutral regions and those with selection strength *s* = 0.01 together, with probabilities around 0.2 *−* 0.3. Regions under stronger selection have much higher probabilities (around 0.9 *−* 1). Although the network was fine-tuned with only a few simulated regions under selection, the real data includes regions of selection, so this general pattern exposes the ability of most discriminators to see selection as realistic.

#### 3.1.2 Validation on known selected regions

To understand how the discriminator behaves when presented with real data from a test population, we produced predictions for regions known to be under selection (from [26]) and compared to predictions from the rest of the genome. We also included data simulated under the demography inferred by the `pg-gan` training run that produced the given discriminator. In theory this is the data which the discriminator thinks is most similar to real data.

Overall we find significant differences in the predictions for selected regions vs. neutral, as seen in the second rows of Figures 3, 4, and 5 (C). For example, in Figure 3, we show the performance of a CEU-trained discriminator (seeds 19) on real data from GBR. Horizontal bars represent means (solid) and 0.05 *−* 0.95 quantiles (dashed). In this case, the t-test p-value for a difference in means between selected (0.423) vs. neutral (0.261) probabilities is 1.63 *×* 10^*−*50^. Of the 50 successfully trained discriminators, the p-value is significant (at the 0.05 confidence level) in all cases. Predictions for simulated data (neutral) and neutral real data typically have very similar distributions, indicating that the generator’s demographic model trained successfully.

For each discriminator, we also performed a genome-wide scan over real test regions (see Figures 3, 4, and 5, E). In the case of CEU (seed 19) for example, we computed predictions for GBR data in 36-SNP regions, then averaged 5 such regions (in overlapping windows) to create the final predictions for these modified Manhattan plots. Similarly, we ran CHS through the CHB-trained discriminators (seed 9 shown) and ESN through the YRI-trained discriminators (seed 4 shown).

#### 3.1.3 Interpretability analysis

After performing correlations between the hidden units of each discriminator and traditional summary statistics (both calculated on the test data), we find some common patterns. Our hierarchical clustering approach shows extensive redundancy in the hidden units, which is to be expected given the dropout computations included in the last fully connected layers. We note that none of the correlation values are particularly high – the largest (in magnitude) are around *±*0.4. Frequently the highest correlations are with rare variants (first entries of the SFS), LD statistics, and *π* (pairwise heterozygosity). For some discriminators, inter-SNP distances also seem to be important. In Figures 3, 4, and 5 (D) we show example heatmaps of correlation values between hidden values of the last layer and summary statistics.

### 3.2 Ensemble results

For the successful discriminators of each training population, we average the probabilities for each test region to create ensemble results. To smooth the results, we further average predictions over five consecutive 36-SNP region (in overlapping windows). Across populations, the mean probability of selection is 0.25, and approximately 1% of the genome has a probability of selection >50% (1.1% for GBR, 1.3% for CHS, and 0.45% for ESN; Figure 6). We confidently identify well-known targets of selection, including *LCT* (probability of selection, p=0.79) [38], *SLC24A5* (p=0.94) [39], *KITLG* (p=0.79) [40, 41] and *OCA2/HERC2* (p=0.86) [42] in GBR ; *EDAR* (p=0.55) in CHS [43]; and *APOL1* (p=0.83) [44] in ESN. We also assign high probabilities of selection to several potentially novel targets, including *APPBP2/PPM1D* (p=0.92) in GBR; *CENPW* (p=0.95) and *EXOC6B* (p=0.95) in CHS; and *EHBP1* (p=0.88), *WHSC1L1* (p=0.87), *ARID1A* (p=0.85) in ESN (Figure 7). Full predictions are given in Supplementary Table S1.

**Figure 6.**
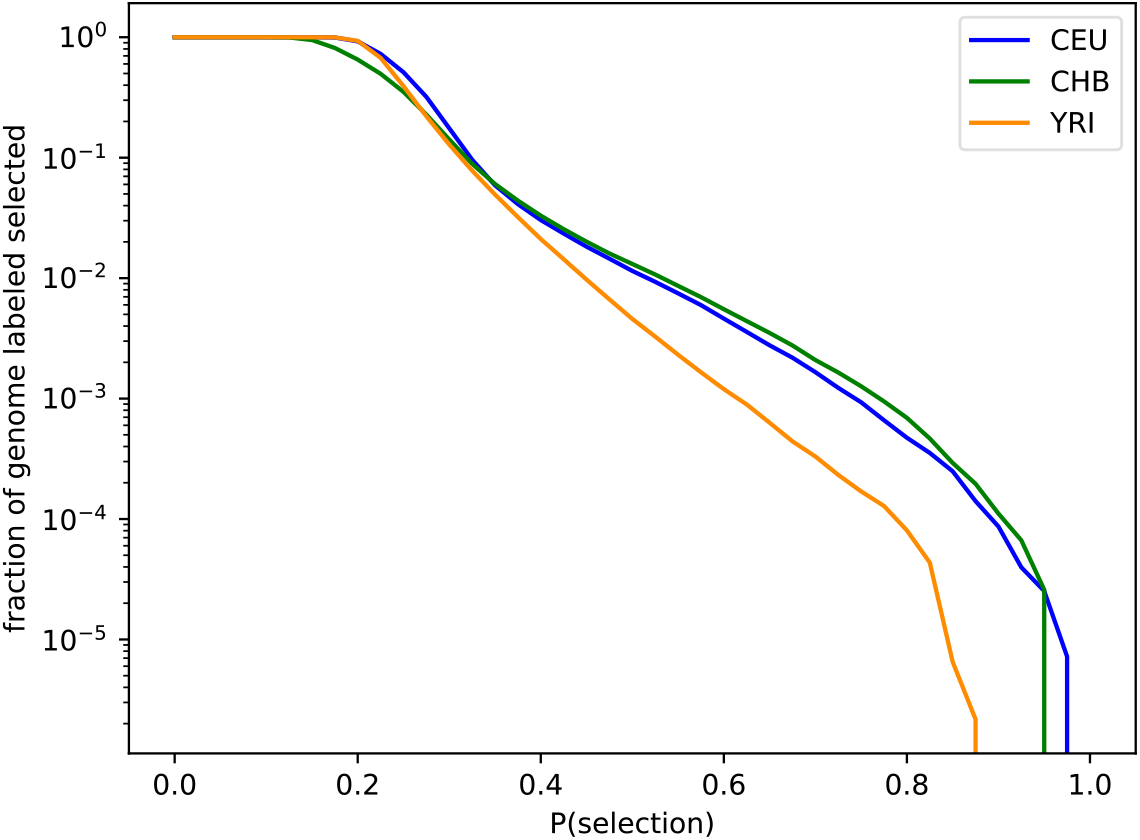
Ensemble probability results. On the x-axis is the probability of selection, and the y-axis shows the fraction of the genome classified as selected if we used the given probability threshold (log scale). Across populations the average probability of selection is 0.25, and approximately 1% of the genome has a probability of seleciton >50%.

**Figure 7.**
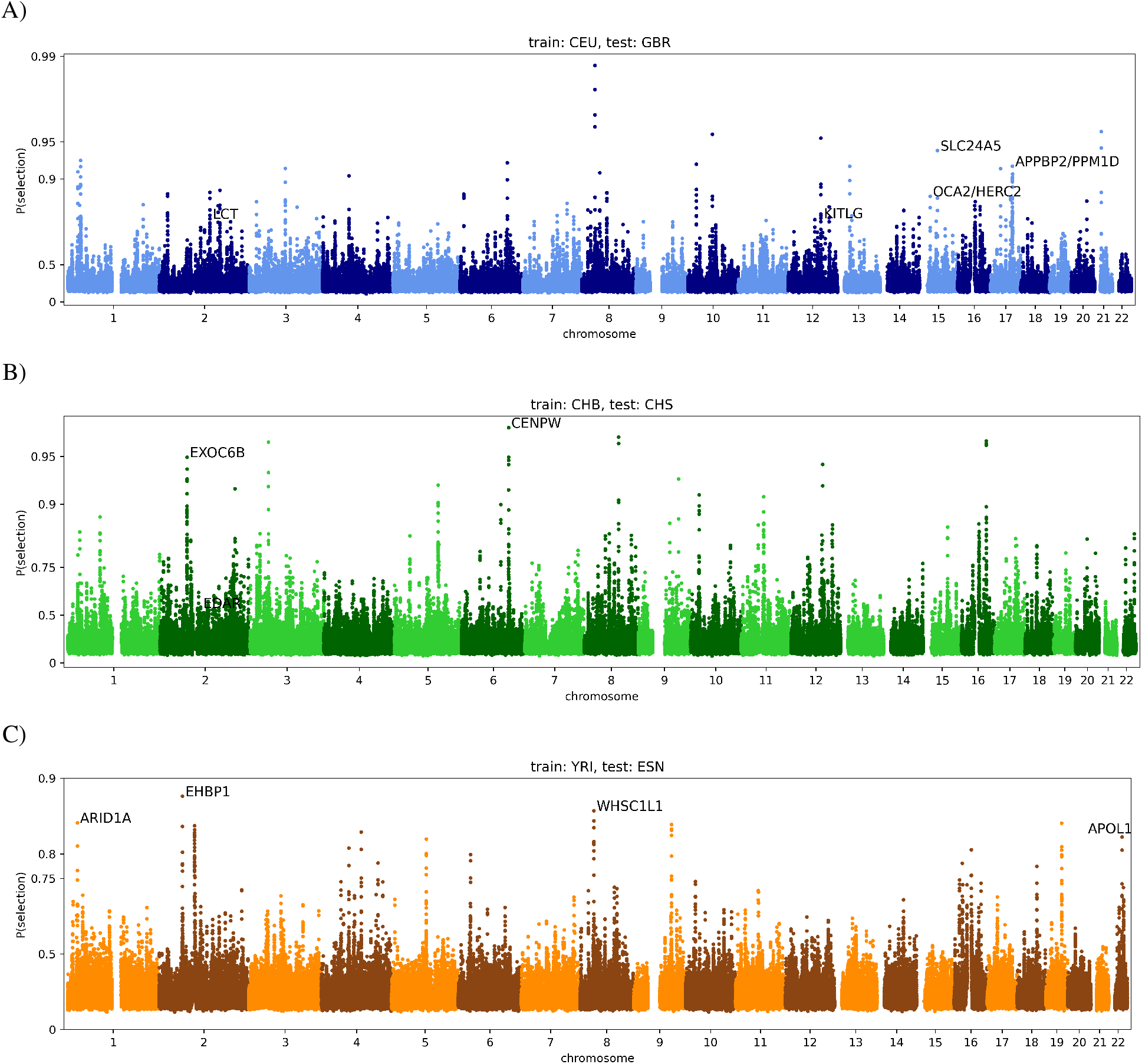
Ensemble results. Genome-wide selections scans for CEU/GBR (A), CHB/CHS (B), and YRI/ESN (C). In each case, the x-axis represents genomic position, the y-axis represents the probability of selection (log scaled) and each point represents the average of five consecutive 36-SNP windows. In black text we highlight known and novel regions with a high probability of selection.

To more systematically validate our results, we obtained selected regions from Grossman *et al* [26] (Supplementary Figure S3), which uses a composite of multiple classical selection statistics to identify selected regions. We calculated our mean prediction probabilities inside and outside these regions and then performed permutation testing by shuffling the locations of the selected regions from [26] randomly throughout the genome. For each of 100 such permutations we re-computed the mean probability of our regions overlapping with these permuted regions. In all cases the mean selection probability of the true known selected genes (*D*(*Y*_*s*_)) was much higher than the permutated values. p-values were 0.0 for all populations and our mean was 8-13 standard deviations above the permutation testing mean (Table 3).

**Table 3:** Ensemble results. For each population we report the mean predicted probabilities for regions that are selected vs. neutral (according to [26]), and the p-value for the difference in mean probabilities between these two sets of regions.

|  | CEU/GBR | CHB/CHS | YRI/ESN |
| --- | --- | --- | --- |
| # successful discriminators | 17/20 | 18/20 | 15/20 |
| mean neutral $D(Y_n)$ | 0.262 | 0.237 | 0.252 |
| mean selection $D(Y_s)$ | 0.386 | 0.385 | 0.350 |
| p-value for difference in means | $8.87 \times 10^{-47}$ | $2.26 \times 10^{-84}$ | $4.92 \times 10^{-83}$ |
| # std dev above mean (permutation testing) | 8.75 | 9.47 | 12.61 |

### 3.3 Runtime analysis

Overall the runtime of our method is dominated by the initial training of `pg-gan`, which takes 5-7 hours per discriminator. After the initial training, fine-tuning is very fast, about 1 minute per discriminator. Selection simulations with `SLiM` were completed in advance of fine-tuning, and were highly sensitive to the demography used to mirror each population. YRI simulations took about 4-5 hours per 100 regions, CHB simulations took about 1 hour per 100 regions, and CEU simulations took about 4-8 hours per 100 regions, depending on the strength of selection (all simulations were parallelized). We hypothesize that CEU took the longest due to the highest inferred exponential growth rate. CHB had the lowest average effective population size, likely resulting in faster simulations.

## 4 Discussion

This work develops three novel aspects of machine learning in population genetics. First, we show how a discriminator trained as part of a GAN can be used as a classifier independent of the generator. Second, we show how the discriminator can be further incentivized to give high probabilities to selected regions in particular through fine-tuning on a small number of selection simulations. Finally, we show how the hidden units of the discriminator can be interpreted in terms of known summary statistics. In particular, we find that rare variant counts, LD statistics, and pairwise heterozygosity are implicitly computed by hidden units in the discriminator network.

A major advantage of our approach is that we do not need to simulate large numbers of selected regions across a high-dimensional space of selection parameters. Training `pg-gan` is not instantaneous, but is much faster than it would be if we used selection simulations throughout the entire training procedure. A typical training run of `pg-gan` uses around 1.5 million neutral simulated regions depending on the number of demographic parameters. In contrast, we only use around five thousand selection simulations for fine-tuning. By adding this step, we are able to quickly create discriminators that have the ability to pick up on real selected regions.

In order to validate our approach in other species, it would be helpful to identify a set of regions under positive selection (for testing). This could be bypassed with high-quality selection simulations, but these might be difficult to obtain depending on the species. Here, we used known selected regions to validate our approach but an alternative would be to instead incorporate these into `pg-gan` training as an additional fine-tuning step. Currently, our interpretability work is on a global scale, meaning that we analyze genome-wide patterns. Future work could examine why the discriminator makes predictions for particular regions, especially outliers with high probabilities.

A caveat of our approach is that the non-neutral regions identified by the discriminator cannot be directly interpreted in terms of selection parameters. Simulations suggest that we can easily detect hard sweeps with selection coefficients greater than around 1%, but may not be able to detect weaker selection (similar to state-of-the art population genetic methods for detecting selection [41, 45]). It is also possible that, despite the fine-tuning step, the regions we identify actually reflect different types of deviation from the generator model, such as heterogeneity in mutation or recombination rates, or structural variation. However, we note that almost all approaches to detecting to selection are sensitive to these effects to some extent and our results, like those of all selection scans, should be treated as candidate regions requiring validation. Another limitation is that we only train and model positive selection. However, our fine-tuning approach could be extended to other selection regimes, for example background and balancing selection and balancing selection. In fact, we could modify the last layer of the discriminator network to output a multi-class prediction that incorporates these different modes of selection. More generally, we view our work as the beginning of an exploration of how the trained discriminator can be used in transfer learning approaches – for these and other evolutionary applications.

## Data availability statement

All human genomic data used in this study is publicly available [35]. Our software is also publicly available open-source at https://github.com/mathiesonlab/disc-pg-gan.

## Acknowledgements

SM is funded in part by NIH grant R15HG011528. The content is solely the responsibility of the authors and does not necessarily represent the official views of the National Institutes of Health.

