## Supplemental Material for "Interpreting Generative Adversarial Networks to Infer Natural Selection from Genetic Data"

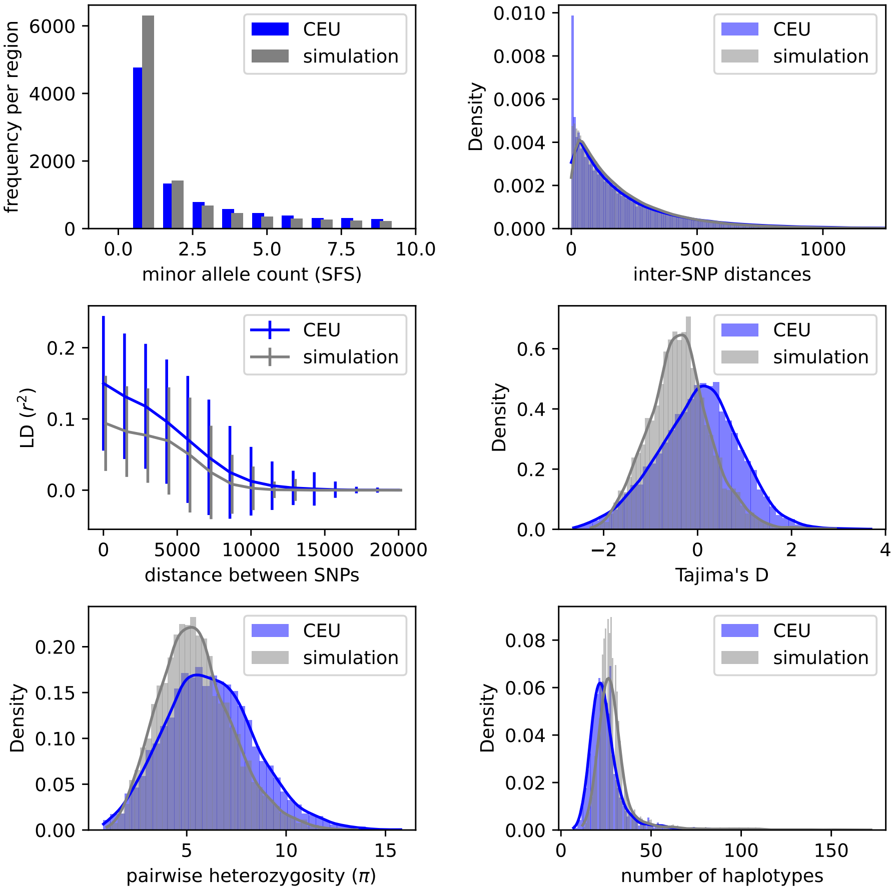


Figure S1: Summary statistic comparison for a failed `pg-gan` training run. In this example (seed 12), CEU was used as the training data and the learning process failed (see Figure S2 below). Typically with failed training, the generator takes a random walk around the parameter space, since the discriminator essentially ignores the input data (whether real or simulated). In most cases this results in an inferred demographic model that fails to recapitulate properties of the real data, which is what we see in these summary statistics.

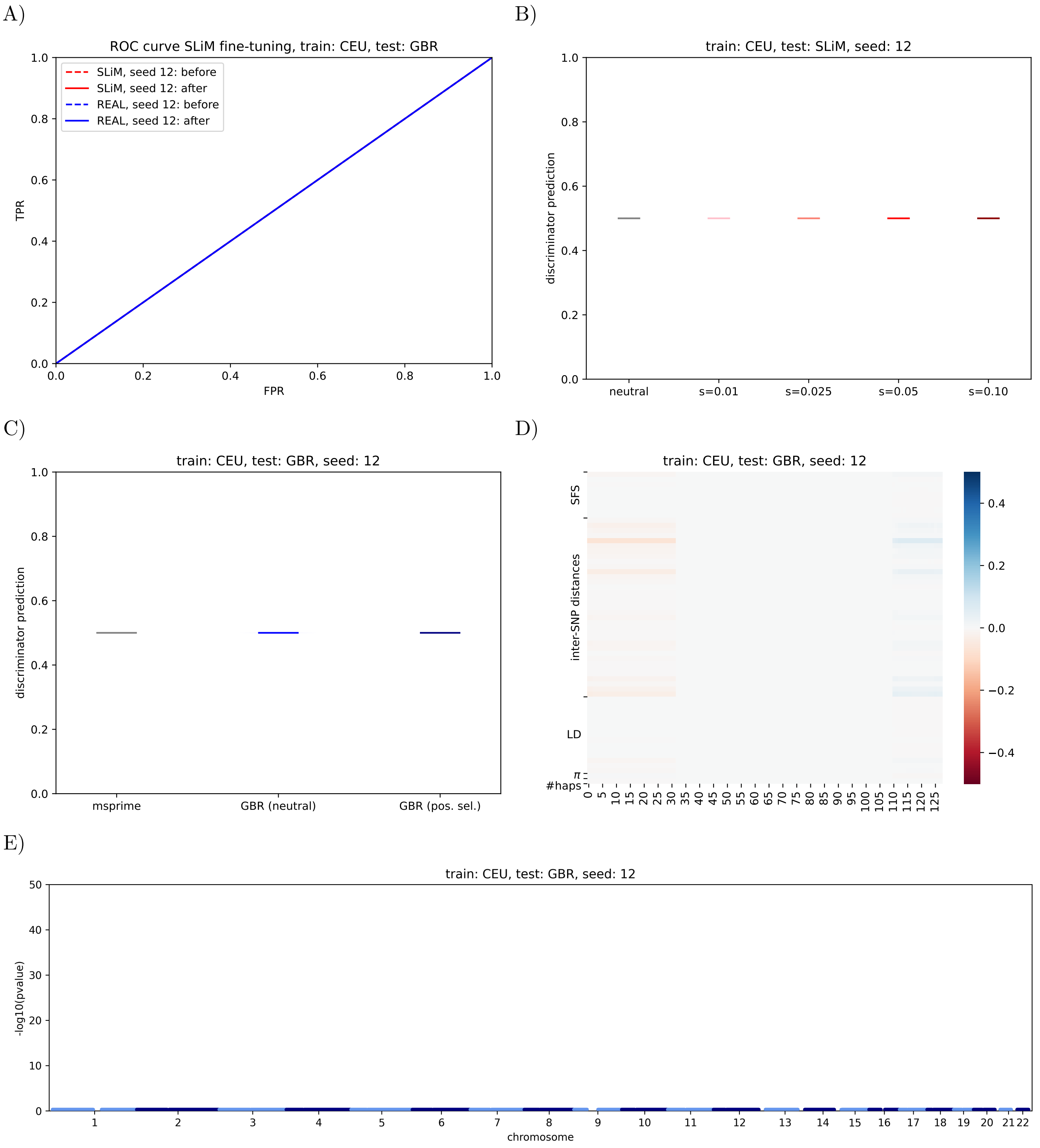


Figure S2: Example of a CEU-trained discriminator with test data from GBR (seed 12). Ten out of the 60 training runs of `pg-gan` failed and show this general pattern, where the discriminator predicts the same value for all regions (essentially ignoring the input data). A) Fine-tuning with selection simulations did not change the random guessing pattern. Thus for the violin plots (B,C), all predictions are the same regardless of the selection coefficient or simulated/real status of the region. D) In our interpretability analysis there is little correlation between discriminator hidden units and any known summary statistics, and no regions are outliers in terms of their probabilities (E).

A)

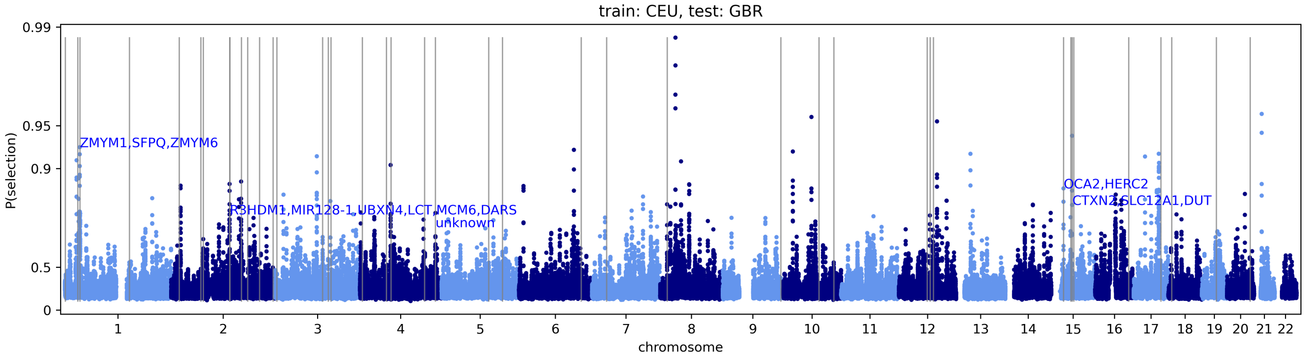


B)

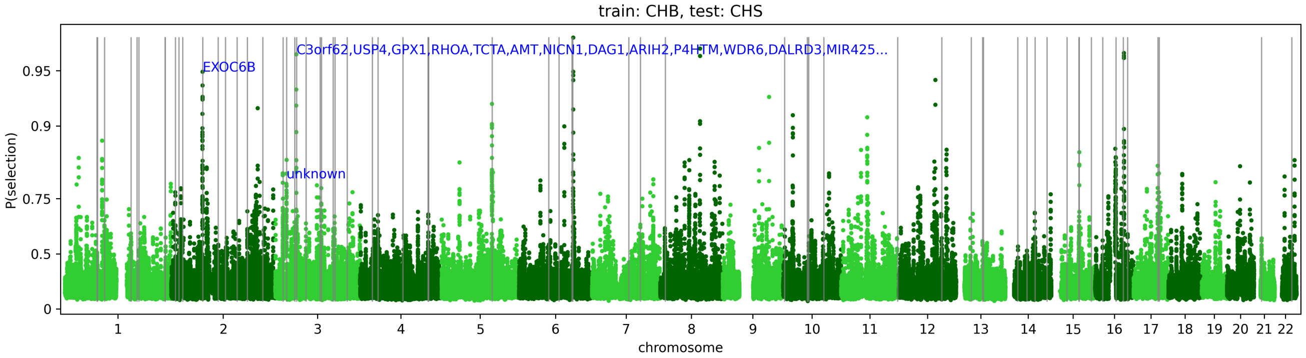


C)

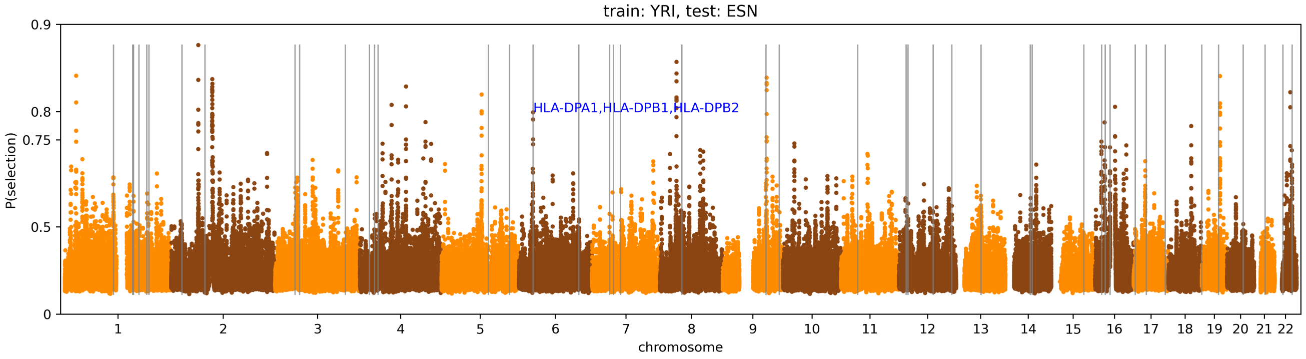


Figure S3: Ensemble results with known selected regions highlighted. Genome-wide selections scans for CEU/GBR (A), CHB/CHS (B), and YRI/ESN (C). In each case, the x-axis represents genomic position, the y-axis represents the probability of selection (plotted on a log scale), each point represents the average of five consecutive 36-SNP windows, and grey vertical lines represent known selected regions from Grossman *et al* (2013). In blue text we highlight regions with P(selection) above 0.75 that also overlap with these known selected regions
